# Plant Cysteine Oxidases are Dioxygenases that Directly Enable Arginyl Transferase-Catalyzed Arginylation of N-End Rule Targets

**DOI:** 10.1101/069336

**Authors:** Mark D. White, Maria Klecker, Richard J. Hopkinson, Daan Weits, Carolin Mueller, Christin Naumann, Rebecca O’Neill, James Wickens, Jiayu Yang, Jonathan C. Brooks-Bartlett, Elspeth F. Garman, Tom N. Grossman, Nico Dissmeyer, Emily Flashman

**Affiliations:** Chemistry Research Laboratory, University of Oxford, 12 Mansfield Road, Oxford, OX1 3TA, United Kingdom; Independent Junior Research Group on Protein Recognition and Degradation Leibniz Institute of Plant Biochemistry (IPB), Weinberg 3, D-06120 Halle (Saale), Germany; ScienceCampus Halle – Plant-based Bioeconomy, Betty-Heimann-Str. 3, D-06120 Halle (Saale), Germany; Institute of Biology I, RWTH Aachen University, Worringerweg 1, D-52074 Aachen, Germany; Chemical Genomics Centre of the Max Planck Society, Department of Chemistry and 19 Chemical Biology, Technische Universität Dortmund, Otto-Hahn-Str. 15, D-44227 Dortmund, Germany; VU University Amsterdam, De Boelelaan 1083, 1081 HV Amsterdam, The Netherlands; Department of Biochemistry, University of Oxford, South Parks Road, Oxford, OX1 3QU, United Kingdom

**Author notes:** **Abbreviations** PCO, plant cysteine oxidase; ATE, arginyl tRNA transferase; ERF-VII, group VII ETHYLENE RESPONSE FACTOR; 2OG, 2-oxoglutarate; NMR, nuclear magnetic resonance; Met, methionine; NME, N-terminal Met excision; Nt, N-terminal; NO, nitric oxide; HIF, hypoxia-inducible factor; PHD, prolyl hydroxylase; MALDI-MS, matrix-assisted laser desorption/ionization-mass spectrometry; LC-MS, liquid chromatography-mass spectrometry; HRE, HYPOXIA RESPONSIVE ERF; RAP, RELATED TO APETALA2; EBP, ETHYLENE RESPONSE FACTOR 72; CDO, cysteine dioxygenase; MAP, Met-aminopeptidase.

**Keywords:** cysteine dioxygenase, ethylene response factors, plant hypoxia, N-end rule pathway, arginyl transferase, E3 ubiquitin ligase, proteostasis

## Abstract

Crop yield loss due to flooding is a threat to food security. Submergence-induced hypoxia in plants results in stabilisation of group VII ETHYLENE RESPONSE FACTORS (ERF-VIIs), which aid survival under these adverse conditions. ERF-VII stability is controlled by the N-end rule pathway, which proposes that ERF-VII N-terminal cysteine oxidation in normoxia enables arginylation followed by proteasomal degradation. The PLANT CYSTEINE OXIDASEs (PCOs) have been identified as catalysts of this oxidation. ERF-VII stabilisation in hypoxia presumably arises from reduced PCO activity. We directly demonstrate that PCO dioxygenase activity produces Cys-sulfinic acid at the N-terminus of an ERF-VII peptide, which then undergoes efficient arginylation by an arginyl transferase (ATE1). This is the first molecular evidence showing N-terminal Cys-sulfinic acid formation and arginylation by N-end rule pathway components, and the first ATE1 substrate in plants. The PCOs and ATE1 may be viable intervention targets to stabilise N-end rule substrates, including ERF-VIIs to enhance submergence tolerance in agronomy.

## Introduction

All aerobic organisms require homeostatic mechanisms to ensure O_2_ supply and demand are balanced. When supply is reduced (hypoxia), a hypoxic response is required to decrease demand and/or improve supply. In animals, this well characterized response is mediated by the Hypoxia-Inducible transcription Factor (HIF), which upregulates genes encoding for vascular endothelial growth factor, erythropoietin and glycolytic enzymes amongst many others.^1–3^ Hypoxia in plants is typically a consequence of reduced O_2_ diffusion under conditions of waterlogging or submergence, or inside of organs such as seeds, embryos, or floral meristems in buds where the various external cell layers act as diffusion barriers. Although plants can survive temporary periods of hypoxia, flooding negatively impacts on plant growth, and if sustained it can result in plant damage or death^4^. This has a major impact on crop yield; for example, flooding resulted in crop loss costing $3 billion in the U.S. in 2011.^5^ As climate change results in increased severe weather events including flooding^4^, strategies to address crop survival under hypoxic stress are needed to meet the needs of a growing worldwide population.

The response to hypoxia in rice, Arabidopsis, and barley is known to be mediated by the group VII ETHYLENE RESPONSE FACTORs (ERF-VIIs).^6–11^ It has been found that these transcription factors promote the expression of core hypoxia-responsive genes, including those encoding alcohol dehydrogenase and pyruvate decarboxylase that facilitate anaerobic metabolism.^12,13^ Crucially, it was shown, initially in Arabidopsis, that the stability of the ERF-VIIs is regulated in an O_2_-dependent manner via the Arg/Cys branch of the N-end rule pathway, which directs proteins for proteasomal degradation depending on the identity of their N-terminal amino acid.^14–16^ Thus, a connection between O_2_ availability and the plant hypoxic response was identified.^11,17,18^ The Arabidopsis ERF-VIIs are translated with the conserved N-terminal motif MCGGAI/VSDY/F^4^ and co-translational N-terminal methionine excision, catalyzed by Met amino peptidases (MAPs)^19,20^, leaves an exposed N-terminal Cys which is susceptible to oxidation.^14–16^ N-terminally oxidized Cys residues (Cys-sulfinic acid or Cys-sulfonic acid, **Supplementary Figure 1**) are then proposed to render the ERF-VII N-termini substrates for arginyl tRNA transferase (ATE)-catalyzed arginylation. The subsequent Nt-Arg-ERF-VIIs are candidates for ubiquitination by the E3 ligase PROTEOLYSIS6 (PRT6)^21^ which promotes targeted degradation via the 26S proteasome. It has also been shown that degradation of ERF-VIIs by the N-end rule pathway can be influenced by NO, and that the ERF-VIIs play a role in plant NO-mediated stress responses.^22,23^

The plant hypoxic response mimics the equivalent well-characterized regulatory system in animals, whereby adaptation to hypoxia is mediated by HIF. In normoxic conditions, HIF is hydroxylated at specific prolyl residues targeting it for binding to the von Hippel-Lindau tumour suppressor protein (pVHL), the recognition component of the E3-ubiquitin ligase complex, which results in HIF ubiquitination and proteasomal degradation.^1,3^ Thus, while not substrates for the N-end rule pathway of protein degradation, HIF levels are regulated by post-translational modification resulting in ubiquitination, in a manner that is sensitive to hypoxia. HIF prolyl hydroxylation is catalyzed by O_2_-dependent enzymes, the HIF prolyl hydroxylases (PHDs 1-3),^2^ which are highly sensitive to O_2_ availability.^24,25^ These O_2_-sensing enzymes are thus the direct link between O_2_ availability and the hypoxic response.^26,27^

Crucially, a family of five enzymes, the PLANT CYSTEINE OXIDASEs (PCO1-5) were identified in Arabidopsis^28^ that were reported to catalyze the O_2_-dependent reaction in the plant hypoxic response, specifically the oxidation of the conserved Cys residue at the N-terminus of the Arabidopsis ERF-VIIs, RAP2.2, RAP2.12, RAP2.3, HRE1 and HRE2. It was found that overexpression of PCO1 and 2*in planta* specifically led to depleted RAP2.12 protein levels and reduced submergence tolerance, while *pcol pco2* T-DNA insertion mutants accumulated RAP2.12 protein. Isolated recombinant PCO1 and PCO2 were shown to consume O_2_ in the presence of pentameric peptides CGGAI corresponding to the N-termini of various ERF-VIIs (**Supplementary Table 1**^28^). The identification of these enzymes indicates that the hypoxic response in plants is enzymatically regulated,^28^ potentially in a similar manner to the regulation of the hypoxic response in animals by the HIF hydroxylases. The PCOs may therefore act as plant O_2_ sensors.

Validation of the chemical steps in the Arg/Cys branch of the N-end rule pathway is still limited, both in animals and plants. We therefore sought to provide molecular evidence that the PCOs catalyze the oxidation step in ERF-VII proteasomal targeting and to determine if this step is required for further molecular priming by arginylation. Using mass spectrometry and NMR techniques we confirm that PCO1 and also PCO4-representatives of the 2 different PCO ‘subclasses’ based on sequence identity and expression behavior^28^ – catalyze dioxygenation of the N-terminal Cys of Arabidopsis ERF-VII peptide sequences to Cys-sulfinic acid (CysO_2_). This oxidation directly incorporates molecular O_2_. To our knowledge, these are the first described enzymes that catalyze cysteinyl oxidation, as well as being the first described cysteine dioxygenases in plants. We then verify that the Cys-sulfinic acid product of the PCO-catalyzed reactions is a direct substrate for the arginyl tRNA transferase ATE1, demonstrating that PCO activity is relevant and sufficient for the subsequent step of molecular recognition and modification according to the N-end rule pathway. This provides the first molecular evidence that Nt-Cys-sulfinic acid is a *bona fide* substrate for N-end rule mediated arginylation. Overall, we thus define the PCOs as plant cysteinyl dioxygenases and ATE1 as an active arginyl transferase, establishing for the first time a direct link between molecular O_2_, PCO catalysis and ATE1 recognition and modification of N-end rule substrates.

## Results

### PCOs catalyze modification of RAP2_2-11_ in an O_2_-dependent manner

N-terminally hexahistidine-tagged recombinant PCO1 and 4 were purified to ~90% purity, as judged by SDS-PAGE **(Supplementary Figure 2a)**. Protein identity was confirmed by comparison of observed and predicted mass by LC-MS (PCO1 predicted mass 36,510 Da, observed mass 36,513 Da; PCO4 predicted mass 30,680 Da, observed mass 30,681 Da, **Supplementary Figure 2b)**. Both PCO1 and PCO4 were found to be monomeric in solution and to co-purify with substoichiometric levels of Fe(II) (~0.3 atoms Fe(II) per monomer, **Supplementary Figure 2c-e**), in line with the reported parameters of recombinant forms of their distant homologs, the cysteine dioxygenases (CDOs).^28–30^ The activity of the purified PCO1 and PCO4 was tested towards a synthetic 10-mer peptide corresponding to the methionine excised N-termini of the ERF-VIIs RAP2.2, RAP2.12, and HRE2 (H2N-CGGAIISDFI-COOH, hereafter termed RAP2_2.11_ **Supplementary Table 1**). Assays comprising RAP_2-11_ at 100 μM in the presence or absence of PCO1 or PCO4 at 0.5 μM underwent aerobic or anaerobic coincubation for 30 minutes at 30°C prior to analysis of the peptide by matrix-assisted laser desorption/ionization-mass spectrometry (MALDI-MS, **Figure 1a,b**;). Only under aerobic conditions and in the presence of PCO1 or PCO4, did the spectra reveal the appearance of two species with mass increases of +32 Da and +48 Da, corresponding to two or three added O atoms, suggesting an O_2_-dependent reaction for PCOs 1 and 4 (**Figure 1b**), as previously shown for PCOs 1 and 2 (note that supplementation of Fe(II) and/or addition of ascorbate was not required for the end-point PCO1/4 activity assays conducted in this study).^28^ These mass shifts were deemed to be consistent with enzymatic formation of Cys-sulfinic (CysO_2_, +32 Da) and Cys-sulfonic acid (CysO_3_, +48 Da; **Supplementary Figure 1**). Although homology between the PCOs and CDOs^28,30^ leads to the predisposition that they will perform similar chemistry (i.e. catalyse Cys sulfinic acid formation), both Cys-sulfinic and Cys-sulfonic acid are proposed to be Arg transferase substrates in the Arg/Cys branch of N-end rule mediated protein degradation and therefore both were considered as potential products of the PCO-catalysed reaction.^14–16^

**Figure 1.**
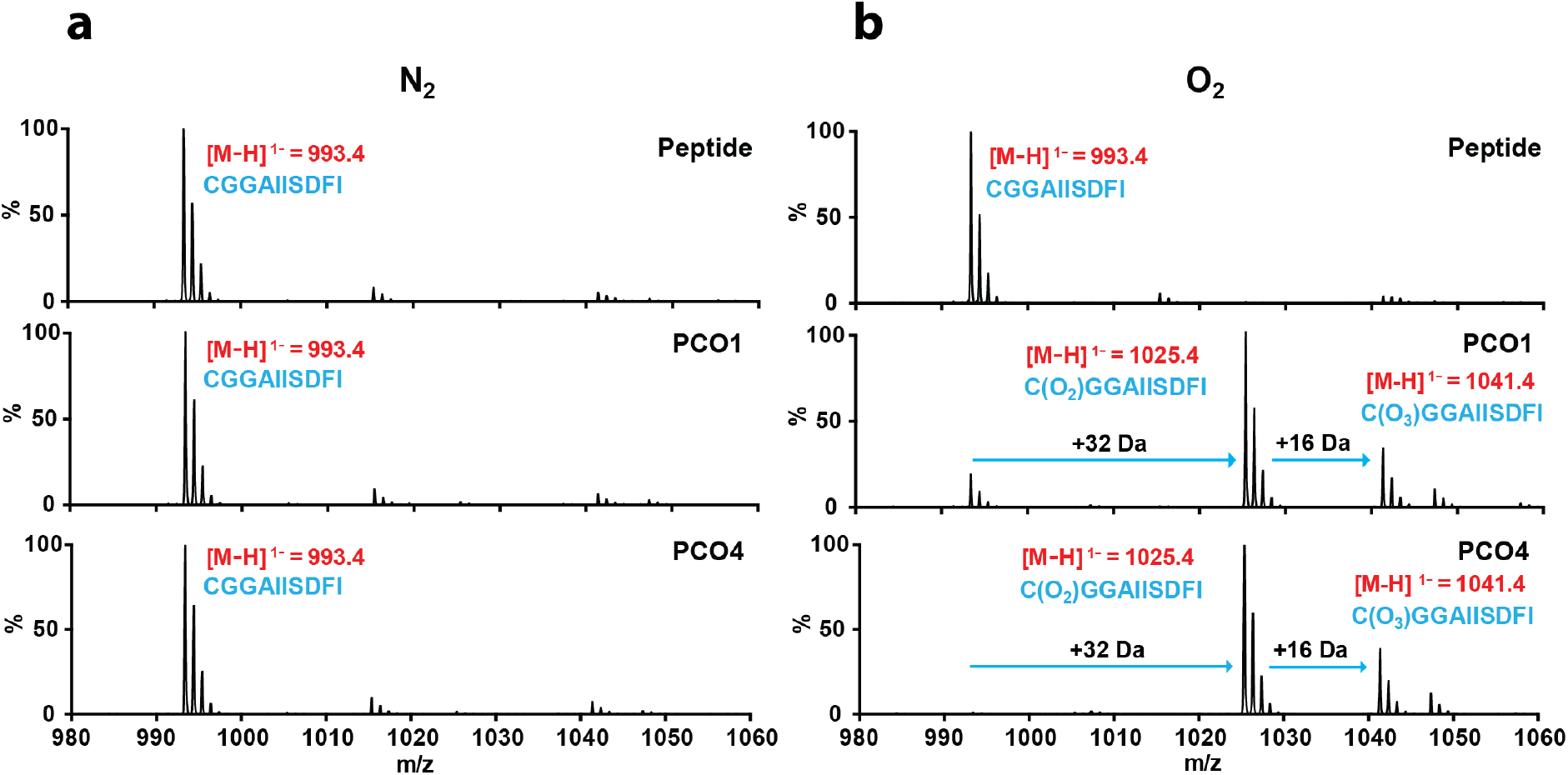
PCO1, PCO4 and O_2_-dependent modification of a RAP2_2__n peptide substrate, consistent with Cys-oxidation.

MALDI-MS spectra showing products following PCO1 and PCO4 incubation with RAP2_2-11_ under anaerobic **(a)** or aerobic **(b)** conditions. Products with mass increases of +32 Da and +48 Da were only observed in the presence of PCO1 or PCO4 and O_2_.

### PCOs are dioxygenases catalyzing the incorporation of both atoms of O_2_ into RAP2_2-11_

To ascertain whether the PCOs function as dioxygenases and thus to confirm a direct connection between molecular O_2_ and PCO activity, we sought to verify the source of the O atoms in the oxidized RAP2_2-11_ by conducting assays in the presence of ^18^O_2_ as the cosubstrate or H2^18^O as the solvent. To probe O_2_ as the source of O atoms in the product, anaerobic solutions of RAP2_2-11_ were prepared in sealed vials before addition of PCO4 using a gas-tight syringe. The vials were then purged with ^16^O_2_ or ^18^O_2_ and the reactions were allowed to proceed at 30°C for a subsequent 20 minutes. Upon analysis by MALDI-MS, the mass of the products revealed that molecular O_2_ was incorporated into the Cys-sulfinic acid product (**Figure 2a**). The Cys-sulfinic acid product had a mass of +32 Da in the presence of ^16^O_2_ and +36 Da in the presence of ^18^O_2_, demonstrating addition of two ^18^O atoms and indicating that O_2_ is the source of O atoms in this product. The Cys-sulfonic acid product had a mass of +52 Da in the presence of ^18^O_2_, indicating a third ^18^O atom had not been incorporated into this product. To probe whether the source of the additional mass in the apparent Cys-sulfonic acid product was an O atom derived from water, an equivalent reaction was carried out under aerobic conditions in the presence of H_2_^18^O (H_2_^18^O:H_2_O in a 3:1 ratio). No additional mass was observed in the peak corresponding to the Cys-sulfonic acid, raising the possibility that the +48 Da species observed by MALDI-MS is not enzymatically formed. Importantly, following incubation in the presence of H2^18^O no additional mass was observed in the peak corresponding to Cys-sulfinic acid, confirming that this species is a product of a reaction where molecular O_2_ is a substrate (**Figure 2b**).

**Figure 2.**
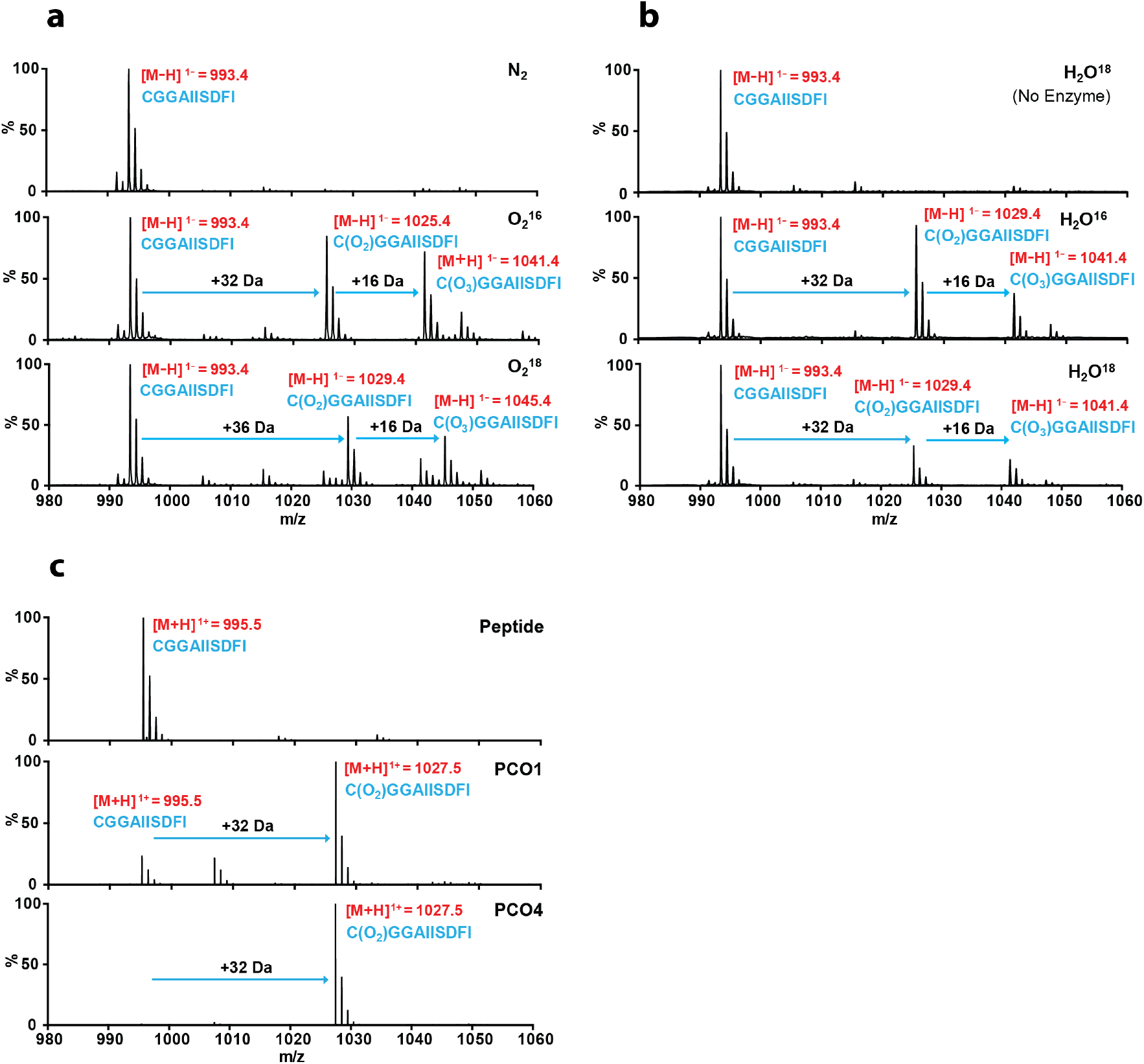
The PCOs are dioxygenases which catalyze incorporation of molecular O_2_ into RAP2_2-11_.

**(a)** MALDI-MS spectra showing that PCO4-catalyzed reactions carried out in the presence of ^18^O_2_ result in a +4 Da increase in the mass of the putative Cys-sulfinic acid product, however a +6 Da increase in the size of the putative Cys-sulfonic acid product is not observed; **(b)** MALDI-MS spectra showing that PCO4-catalyzed reactions carried out in the presence of H2^18^O show no additional incorporation of mass compared with products of reactions in the presence of H_2_^16^O; **(c)** LC-MS spectra confirm that the +48 Da reaction product is an artifact of MALDI-MS analysis **(Supplementary Figure 3)** and incubation of PCO1 and PCO4 with RAP2_2-11_ results in formation of a single product with a mass increase of +32 Da, consistent with Cys-sulfinic acid formation.

To further investigate whether the PCO-catalyzed product species observed at +48 Da is enzymatically produced or an artefact of the MALDI-MS analysis method, we turned to liquid chromatography-mass spectrometry (LC-MS) to analyze the products of the PCO-catalyzed reactions. Under these conditions, only peptidic product with a mass increase of +32 Da was observed after incubation with both PCO1 and PCO4, corresponding to the incorporation of two O atoms and the formation of Cys-sulfinic acid (**Figure 2c**), consistent with the products observed using ^18^O_2_ and H_2_^18^O (**Figure 2a,b**). No product was observed with a mass corresponding to Cys-sulfonic acid, which suggested that the +48 Da product detected by MALDI-MS was indeed an artefact. When combined with the observation that significant quantities of Cys-sulfonic acid were not seen in no-enzyme or in anaerobic controls (**Figure 1**), it was hypothesized that the Cys-sulfinic acid product of the PCO-catalyzed reaction is non-enzymatically converted to Cys-sulfonic acid during MALDI-MS analysis, potentially as a result of laser exposure. Upon subjecting the products of PCO1 and 4 turnover to MALDI-MS analysis with increasing laser intensity, a direct correlation between laser intensity and the ratio of Cys-sulfonic acid:Cys-sulfinic acid product was observed **(Supplementary Figure 3a)**. Of note, significant levels of laser induced formation of +32 and +48 Da species upon analysis of unmodified peptide were not observed **(Supplementary Figure 3b)**. Together, these results confirm that the +48 Da species observed following incubation of the PCOs with RAP2_2-11_ are a product of Cys-sulfinic acid exposure to the MALDI-MS laser, and not a product of the PCO-catalyzed reaction. Overall, these data demonstrate that the PCOs are dioxygenase enzymes, similar to the mammalian and bacterial cysteine dioxygenases (CDOs) to which they show sequence homology.^28,30^

### PCOs catalyze oxidation of N-terminal Cys of RAP2_2-11_ to form Cys-sulfinic acid

Recombinant PCO1 and PCO2 were reported to consume O_2_ in the presence of pentameric CGGAI peptides corresponding to the methionine-excised N-terminus of the Arabidopsis ERF-VIIs.^28^ To definitively verify that the N-terminal cysteinyl residue of RAP2_2-11_ is indeed the target for the PCO-catalyzed +32 Da modifications, we conducted LC-MS/MS analyses on the reaction products. Fragmentation of RAP2_2-11_ that had been incubated in the presence and absence of PCO1 and PCO4 revealed b-and y-ion series consistent with oxidation of the N-terminal Cys residue (**Figure 3a**), confirming that PCOs 1 and 4 act as cysteinyl dioxygenases.

**Figure 3.**
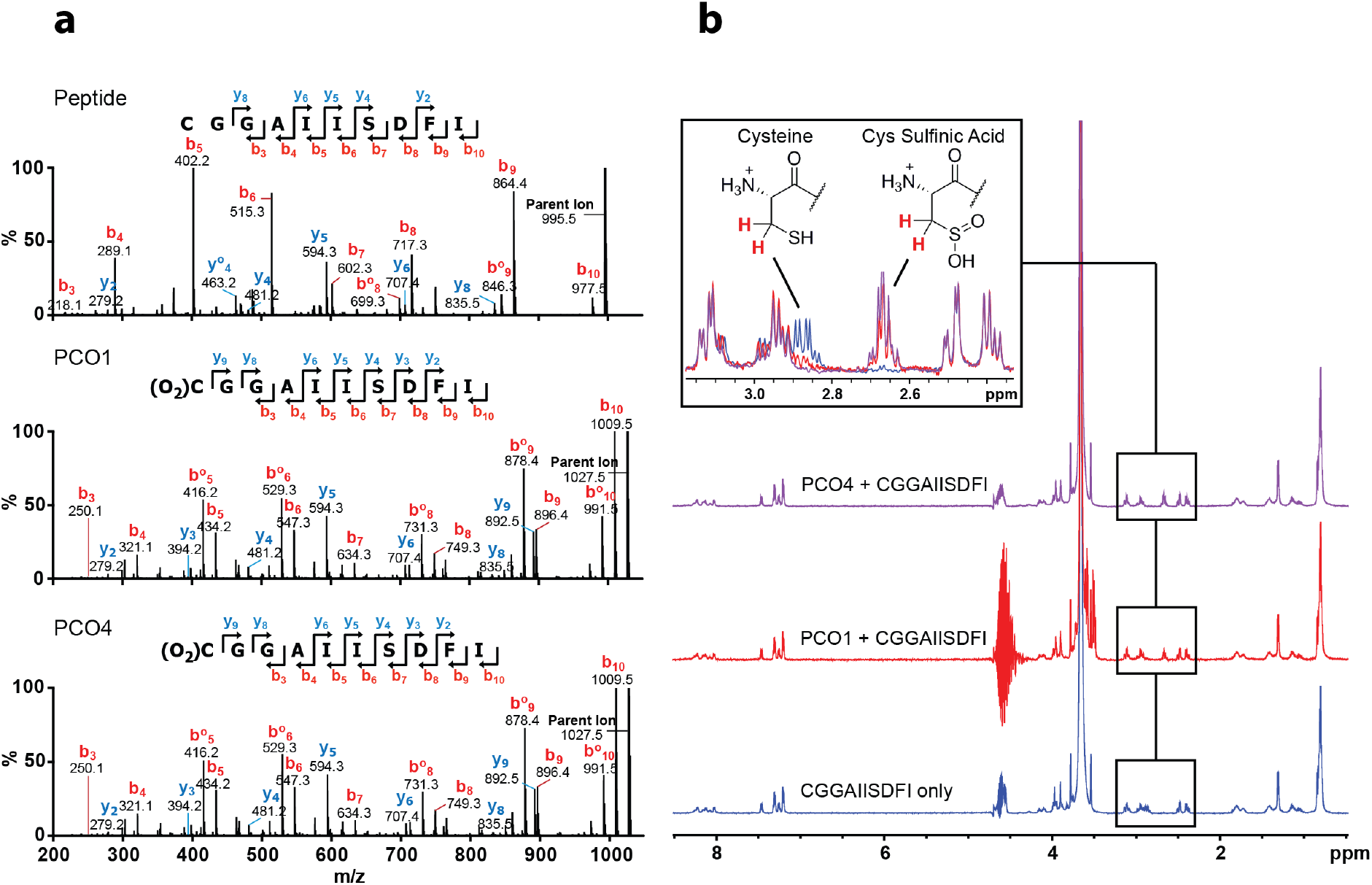
PCO1 and PCO4 oxidize the N-terminal Cys of RAP2_2-11_ to Cys-sulfinic acid as confirmed by (a) LC-MS/MS and (b) ^1^H-NMR.

**(a)** Peptidic products of PCO-catalyzed reactions were subjected to LC-MS/MS analysis. In the presence of enzyme, fragment assignment was consistent with expected b-and y-series ion masses for RAP22-11 with N-terminal Cys-sulfinic acid. **(b)** ^1^H-NMR was used to monitor changes to RAP22-11 (500 μM) upon incubation with enzyme (5 μM). In the presence of PCO1 (red) and PCO4 (purple), the ^1^H-resonance at δ_H_ 2.88 ppm (assigned to the β-cysteinyl protons of RAP2_2-11_, blue) was observed to decrease in intensity, with concomitant emergence of a resonance at δ_H_ 2.67 ppm. This new resonance was assigned to the β-protons of Cys-sulfinic based on chemical shift analysis (see **Supplementary Figure 4**).

As a final confirmation of the nature of the reaction catalyzed by PCO1 and PCO4, their activity was monitored using ^1^H-NMR. Reactions were initiated by adding 5 μM enzyme to 500 μM RAP2_2-11_ (in the presence of 10% D_2_O) and products of the reaction were analysed using a 600 MHz NMR spectrometer. In the presence of both PCO1 and PCO4, modification to the cysteinyl residues was observed, as exemplified by the disappearance of the ^1^H-resonance corresponding to the β-cysteinyl protons (at δ_H_ 2.88 ppm) and the emergence of a new ^1^H-resonance at δ_H_ 2.67 ppm (**Figure 3b**). The chemical shift of the new resonance is similar to that observed for L-Cys conversion to L-Cys-sulfinic acid by mouse CDO,^31^ and also to the chemical shift of an L-Cys-sulfinic acid standard measured under equivalent conditions to the PCO assays **(Supplementary Figure 4)**. Therefore, the resonance shift observed upon PCO1/4 reaction was assigned to the β-protons of L-Cys-sulfinic acid. Overall these results provide confirmation at the molecular level that Arabidopsis PCOs 1 and 4 act as plant cysteinyl dioxygenases, catalyzing incorporation of O_2_ into N-terminal Cys residues on a RAP2 peptide to form Cys-sulfinic acid.

### ATE1 arginylates acidic N-termini including Cys-sulfinic acid

We next sought to confirm that the PCO-catalyzed Cys-oxidation to Cys-sulfinic acid renders a RAP2 peptide capable of and sufficient for onward modification by ATE1. Cys-sulfinic acid has been proposed as a substrate for ATE1 on the basis of its structural homology with known ATE1 substrates Asp and Glu, but evidence has only been reported to date for arginylation of Cys-sulfonic acid.^32,33^ We further sought to validate the role of a plant ATE1: To date ATE1 has been suggested to be responsible for transfer of ^3^H-arginine to bovine α-lactalbumin in highly purified plant extracts *in vitro*^34^ and RAP2.12 stabilization in *ate1 ate2* double null mutant plant lines implicates ATE1 as an ERF-VII-targeting arginyl transferase *in vivo*,^17,18^ To this end, we produced recombinant hexahistidine-tagged Arabidopsis ATE1 **(Supplementary Figure 5)** for use in an arginylation assay which detects incorporation of radiolabeled ^14^C-Arg into biotinylated peptides. C-terminally biotinylated RAP2_2-13_ peptides (H2N-XGGAIISDFIPP(PEG)K(biotin)-NH_2_) where the N-terminal residue, X, constitutes Gly, Asp, Cys or Cys-sulfonic acid were subjected to the arginylation assay in the presence or absence of PCO1/4 (**Figure 4a**). Peptide with an N-terminal Gly did not accept Arg, while an N-terminal Asp did accept Arg, independent of the presence of PCO1 or 4. A peptide comprising an N-terminal Cys-sulfonic acid was also shown to be a substrate for ATE1, again independent of the presence of PCO1 or 4, which is in line with proposed steps of the Arg/Cys N-end rule pathway and has also recently been reported using a similar assay with mouse ATE1.^14–16,35^ Crucially, in the absence of PCO1/4, RAP2_2-13_ with an N-terminal Cys was not an acceptor of arginine transfer by ATE1, yet when either PCO1 or PCO4 was incorporated in the reaction, significant ATE1 transferase activity was observed (**Figure 4a**).

**Figure 4.**
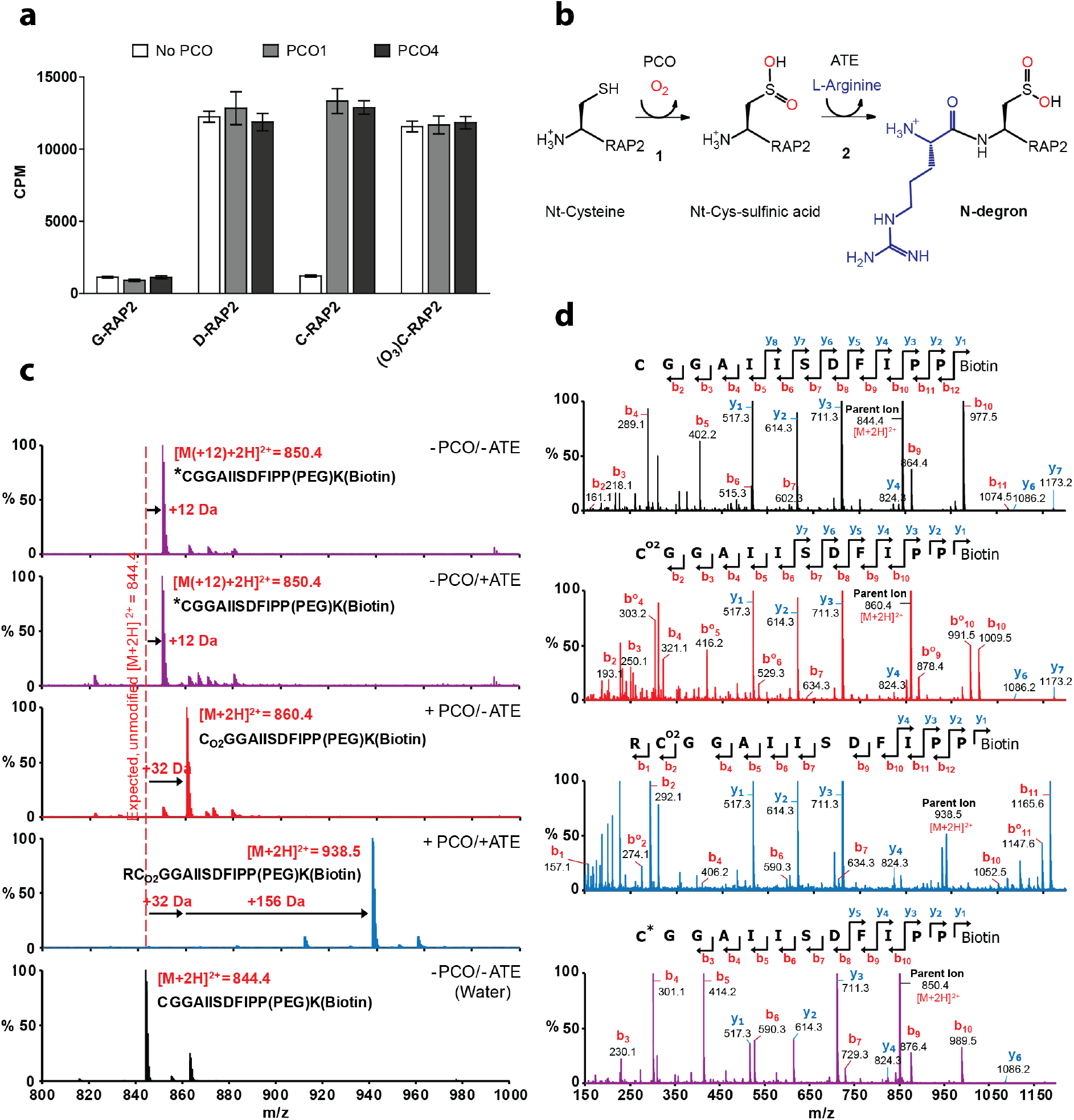
PCO-Catalyzed Cys-sulfinic acid formation renders RAP22-13 a substrate for ATE1 catalyzed arginylation. **(a)** ^14^C-Arg incorporation by ATE1 into the 12-mer N-terminal RAP2 peptide (H<sub>2</sub>N-XGGAIISDFIPP(PEG)K(biotin)-NH<sub>2,</sub> X = Gly, Asp, Cys or Cys-sulfonic acid (C(O<sub>3))),</sub> was assayed by liquid scintillation counting of immobilized biotinylated peptides after the arginylation reaction and removal of unreacted ^14^C-Arg (n=3). In the case of the Cys-starting peptide (RAP2<sub>2-13),</sub> ATE1 activity was strongly dependent on the presence of PCO1 or PCO4. **(b)** Scheme showing PCO-and ATE1-catalyzed reactions on Nt-Cys RAP2.2, as validated in this study. **(c)** LC-MS spectra of products of equivalent assays with Cys-initiated RAP2_2-13_ using non-radiolabelled Arg, revealing a sequential mass increase of +32 (corresponding to oxidation) and +156 Da (corresponding to arginylation) only in the presence of PCO and ATE1 (blue spectrum). The red spectrum shows a +32 Da mass increase for Cys-RAP2_2-1_3 incubated +PCO/-ATE, demonstrating Cys-sulfonic acid formation as expected. Purple spectra show +12 Da products formed upon incubation of Cys-RAP2_2-1_3 in the absence of PCO +/-ATE (for explanation of this mass increase see text and **Supplementary Figure 6**); the black spectrum shows Cys-RAP2_2_”_1_3 dissolved in H_2_O. **(d)** b-and y-ion series spectra generated by MS/MS analysis of Cys-RAP2_2-13_ only (no incubation; black), Cys-RAP22-13 incubated +PCO/-ATE (red), Cys-RAP22-13 incubated with PCO and ATE1 (blue) and Cys-RAP22-13 incubated without PCO or ATE1 (purple), confirming arginylation only at the N-terminus of PCO-modified RAP2_2-13_.

To confirm that the increased detection of radiolabelled arginine corresponded to arginyl incorporation at the N-termini of the peptides, the experiment was repeated using non-radiolabeled arginine in the presence and absence of PCO4 and ATE1, and peptide products subjected to LC-MS analysis (**Figure 4c**). As with RAP2_2-11_ (**Figure 2c**), the Cys-initiated RAP2_2-13_ peptide displayed a +32 Da increase in mass upon incubation with PCO4 only (**Figure 4c**, red spectrum). Importantly, following incubation of Cys-initiated RAP2_2-13_ with both PCO4 and ATE1, a mass increase equivalent to oxidation coupled to arginylation (+188 Da) was observed (**Figure 4c**, blue spectrum). Subsequent tandem MS analysis of these product ions revealed fragmentation species consistent with the assumption that oxidation and sequential arginylation occur at the N-terminus of PCO4-and ATE1-treated peptides (**Figure 4d**, blue spectrum), strongly suggesting that the PCO-oxidized N-termini of ERF-VIIs are rendered N-degrons via additional arginylation (**Figure 4b**).

A +12 Da mass increase was observed in products of control assays lacking PCO4 (**Figure 4c, d**; purple spectra). This appeared to be related to prolonged incubation in the presence of HEPES and DTT as used in the arginylation assay buffer: The +12 Da modification was not observed if the peptide was dissolved in H_2_O (**Figure 4c**, black spectrum) or if incubated with HEPES and DTT for just 1 hour, but was observed when the peptide was incubated with HEPES and DTT overnight **(Supplementary Figure 6)**. It is proposed that under these conditions, trace levels of contaminating formaldehyde react with free Nt-Cys residues to form thiazolidine N-termini.^36^

These results are in line with proposed arginylation requirements for the Arg/Cys branch of the N-end rule pathway^14–16^ including the known Cys-initiated arginylation targets from mammals.^32,33,35,37^ Importantly, these results demonstrate for the first time Arg transfer mediated by a plant ATE dependent on the N-terminal residue of its substrate, and also that both Cys-sulfinic acid (the product of PCO-catalysis) and Cys-sulfonic acid can act as substrates for ATE1. In particular, the arginylation observed with PCO-catalyzed Cys-sulfinic acid supports the assumption that N-terminal residues sterically and electrostatically resembling Asp or Glu can serve as Arg acceptors in reactions catalyzed by ATEs,^33^ and also confirms the importance of the PCOs as a connection between the stability of their ERF-VII substrates and O_2_ availability (**Figure 4b**).

## Discussion

The PCOs were identified in *Arabidopsis thaliana* as a set of five enzymes suggested to catalyze oxidation of N-terminal cysteine residues in ERF-VII transcription factors and oxygen consumption was demonstrated for reactions with short peptides corresponding to their N-termini^28^. This putative oxidation was associated with destabilization of the ERF-VIIs, presumably by rendering them substrates of the Arg/Cys branch of the N-end rule pathway.^14,16^ Under conditions of sufficient O_2_ availability, ERF-VII protein levels are decreased, while under hypoxic conditions, such as those encountered upon plant submergence or in the context of organ development, ERF-VII levels remain high.^17,18^ Importantly, the ERF-VII transcription factors are known to upregulate genes which allow plants to cope with or respond to submergence.^13^ The PCOs are proposed to act as potential O_2_ sensors involved in regulating the plant hypoxic response.^28^

We sought to biochemically confirm the role of the PCOs in the plant hypoxic response, and present here mass spectrometry and NMR data that clearly demonstrate that two enzymes from different ‘subclasses’ of this family, PCOs 1 and 4, are dioxygenases that catalyze direct incorporation of O_2_ into RAP2_2-11_ peptides to form Cys-sulfinic acid. Their direct use of O_2_ supports the proposal that these enzymes may act as plant O_2_ sensors.^28^ A relationship has been demonstrated between O_2_ concentration and PCO activity,^28^ but it will be of interest to perform detailed kinetic characterization of these enzymes to ascertain their level of sensitivity to O_2_ availability, in particular to determine whether their O_2_-sensitivity is similar to that of the HIF hydroxylases in animals.^24,25^ Although there is functional homology between the PCOs and the HIF hydroxylases, they are apparently mechanistically divergent: The PCOs show sequence homology to the Fe(II)-dependent CDO family of enzymes which do not require an external electron donor for O_2_ activation,^28,30^ while the HIF hydroxylases are Fe(II)/2OG-dependent oxygenases. They also co-purified with Fe(II) as reported for both the CDOs^29^ and PHD2^38^. Of note, the PCOs are the first identified CDOs in plants. Further, in contrast to the reactions of mammalian and bacterial CDOs which oxidize free L-Cys, the PCOs are also, to our knowledge, the first identified cysteinyl (as opposed to free L-Cys) dioxygenases.

According to the Arg/Cys branch of the N-end rule pathway, N-terminal Cys oxidation is proposed to enable successive arginylation by ATE1 to render proteins as N-degrons. While both Cys-sulfinic and Cys-sulfonic acid are repeatedly reported as potential arginylation substrates^14–16^, detailed evidence has only been presented to date for arginylation of Cys-sulfonic acid^32,33^ and this only in a mammalian system. We therefore sought to demonstrate that PCO-catalyzed ERF-VII N-terminal Cys oxidation to Cys-sulfinic acid promotes arginylation by ATE1. The arginylation assay and mass spectrometry results we present demonstrate that the PCO-catalyzed dioxygenation reaction is sufficient to trigger N-terminal arginylation of RAP2s by ATE1, thus likely rendering ERF-VIIs (at least those comprising the tested N-terminal sequence) as N-degrons, i.e. recognition by PRT6 and other potential E3 ubiquitin ligases, polyubiquitination and possibly transfer to the 26S proteasome for proteolysis.^14–16^ Collectively therefore, we present the comprehensive molecular evidence confirming the Cys-oxidation and subsequent arginylation steps of the Arg/Cys branch of the N-end rule pathway.^32,33,37^ We also confirm that ATE1 is able to selectively arginylate, as predicted,^33^ acidic N-terminal residues of plant substrates, including Cys-sulfonic acid.

Arginylation has been known as a posttranslational modification since 1963,^39^ to possess a general aminoacyl transferase function in plants (rice and wheat) since 1973^40^ and to have a speculative involvement in the N-end rule pathway since 1988.^41,42^ ATE1 is reported as being capable of arginylating proteins at both acidic N-termini and midchain acidic side chains via canonical and non-canonical peptide bonds, respectively.^43^ Reports of midchain arginylation highlighted a potentially broad involvement of ATEs in posttranslational protein modifications for various functions^35,43,44,45^ but was only very recently brought into question by ^14^C-Arg incorporation assays using arrays of immobilized synthetic peptides.^35,43,44^ To date however only one physiological and two *in vitro* substrates for the Arg/Cys branch of the N-end rule pathway have been characterized, namely mammalian regulator of G protein signaling (RGS) 4, and RGS5 and 10 respectively,^46^ where Nt-Cys oxidation was described (to Cys-sulfonic acid) as was Nt-Cys arginylation.^33,37^ The first non Cys-branch N-end rule arginylation target was shown to require posttranslational proteolytic cleavage of a (pre-)-proprotein. The C-terminal fragment of proteolytically cleaved mouse BRCA1 is Asp-initiated^47^ and gets degraded in an N-end rule-dependent manner. Then, the molecular chaperone BiP (GRP78 and HSPA5, heat shock 70 kDa protein 5) and the oxidoreductase protein disulphide isomerase (PDI), present Glu or Asp after cleavage of their signal peptide, respectively, and were suggested but not shown as putative N-end rule substrates (Hu et al.JBC, 2006). Only very recently, BiP and PDI were identified in mammalian cell culture together with the Glu-initiated calreticulin (CRT) as arginylation targets with a function in autophagy rather than the N-end rule degradation.^48^

Similarly, data regarding the molecular requirements of plant ATEs are limited. Already in 1973, a general aminoacyl transfer activity was found in rice and wheat cell extracts, however, the nature of enzyme, acceptor position and mechanism remained unclear. It was suggested that the N-terminus could serve as Arg acceptor.^40^

The first description of a mutant of the single translatable *ATE1* gene in the Arabidopsis accession Wassilewskija (Ws-0) highlighted a role of ATEs in plant development. Ws-0 lacks the second *bona fide* ATE, that is ATE2, due to a single nucleotide polymorphism in *ATE2* causing a premature stop.^49^ Developmental functions of the single homolog ATE1 in the moss *Physcomitrella patens* were recently described.^50^ Interaction partners of the enzyme were found as well as four arginylated peptides immunologically detected by using antibodies directed against peptides mimicking N-terminal Arg-Asp or Arg-Glu.^51^ In one case, that is the acylamino-acid-releasing enzyme PpAARE, which presents for unknown reasons a neoN-terminal Asp residue which was formerly Asp2 and therefore initiated by Met, an N-terminal arginylation was found with high confidence.Previously, Arg transferase function of Arabidopsis ATE1/2 has been shown using an assay detecting conjugation of ^3^H-Arg to bovine α-lactalbumin (bearing an N-terminal Glu) in the presence of plant extracts from wild type Arabidopsis, and *ate1* and *ate2* single mutants but not from *ate1 ate2* double mutant seedlings.^34^ Therefore, the results we present here demonstrate for the first time Arg transferase activity of a plant ATE towards known plant N-end rule substrates.

Interestingly, in combination with O_2_, nitric oxide was identified as an RGS oxidizing agent, suggesting a potential role of S-nitrosylation in the Arg/Cys branch of the N-end rule pathway, albeit non-enzymatically controlled.^32^ It has also been reported *in planta* that both NO and O_2_ are required for ERF-VII degradation, potentially at the Cys oxidation step.^22,23^ Although in N-end rule-mediated RGS4/5 degradation it has been proposed that Cys-nitrosylation precedes Cys-oxidation (also currently considered a non-enzymatic process), we find that under the conditions used, the PCO1/4-catalyzed reaction does not require either prior cys-nitrosylation or exogenous NO to proceed efficiently. We cannot rule out that NO plays a role in formation of a Cys-sulfonic acid product, which is also a substrate for ATE1 as shown in our Arg transfer experiments. Alternatively, NO may have a role in ERF-VII degradation *in vivo* via non-enzymatic oxidation or via a secondary mechanism. The manner in which NO contributes to Arg/Cys branch of the N-end rule pathway therefore remains to be elucidated.

ERF-VII stabilization has been shown to result in improved submergence tolerance, elegantly demonstrated in barley by mutation of the candidate E3-ubiquitin ligase *PRT6*,^11^ but also in rice containing the *Sub1A* gene; SUB1A is an apparently stable ERF-VII that confers particular flood tolerance in certain rare varieties of rice.^9,17^ Overexpression of *Sub1A* in more commonly grown rice varieties has resulted in a 45% increase in yield relative to *sub1a* mutant lines after exposure to flooding.^52^ If ERF-VII stabilization is indeed a proficient mechanism for enhancing flood tolerance, then manipulation of PCO or ATE activity may be an efficient and effective point of intervention. This work presents molecular validation of their function, providing the basis for future targeted chemical/genetic inhibition of their activity. It also highlights genetic strategies for breeding via introgression of variants of N-end rule pathway components or introduction of alleles of enzymatic components of the N-end rule pathway from non-crop species into crops. Any of these strategies has the potential to result in stabilized ERF-VII levels and therefore increase stress resistance and may therefore help to address food security challenges.

## Online Methods

### Peptide Synthesis, Purification and Characterization

All reagents used were purchased from Sigma-Aldrich unless otherwise stated. The 10-mer RAP2_2-11_ peptide (H_2_N-CGGAIISDFI-COOH) was purchased from GL Biochem (Shanghai) Ltd, China **(Supplementary Table 1)**. The sequence of the 12-mer peptides used in the coupled oxidation-arginylation assay is derived from RAP2.2 and RAP2.12 (H_2_N-X-GGAIISDFIPP(PEG)K(biotin)-NH_2_) and synthesized by Fmoc-based solid-phase peptide synthesis (SPPS) on NovaSyn®TGR resin (Merck KGaA, **Supplementary Table 2**). Fmoc protected amino acids (Iris Biotech GmbH) were coupled using 4 equivalents of (eq) of the amino acid according to the initial loading of the resin. 4 eq amino acid was mixed with 4 eq O-(6-chlorobenzotriazol-1-yl)-N,N,N',N'-tetramethyluronium hexafluorophosphate (HCTU) and 8 eq N,N-diisopropylethylamine (DIPEA; Santa Cruz Biotechnology, sc-293894) and added to the resin for 1 h. In a second coupling, the resin was treated with 4 eq of the Fmoc-protected amino acid mixed with 4 eq benzotriazole-1-yl-oxy-tris-pyrrolidino-phosphonium hexafluoro-phosphate (PyBOP) and 8 eq 4-methylmorpholine (NMM) for 1 h. After double coupling a capping step to block free amines was performed using acetanhydride and DIPEA in N-methyl-2-pyrrolidinone (NMP) (1:1:10) for 5 min. The *C*-terminal Fmoc-Lys(biotin)-OH, the 8-(9-fluorenylmethyloxycarbonyl-amino)-3.6-dioxaoctanoic acid (PEG) linker and the different Fmoc protected N-terminal amino acids were coupled manually. The remaining peptide sequence was assembled using an automated synthesizer (Syro II, MultiSynTech GmbH). Fmoc deprotection was performed using 20 % piperidine in dimethylformamide (DMF) for 5 min, twice. After each step the resin was washed 5 times with DMF, DCM and DMF, respectively. Final cleavage was performed with 94 % trifluoroacetic acid (TFA), 2.5 % 1,2-ethanedithiole (EDT) and 1 % triisopropylsilane (TIPS) in aqueous solution for 2 h, twice. The cleavage solutions were combined and peptides were precipitated with diethyl ether (Et_2_O) at −20°C for 30 min. Peptides were solved in water/acetonitrile (ACN) 7:3 and purified by reversed-phase high performance liquid chromatography (HPLC; Nucleodur C18 culumn; 10×125 mm, 110 Å, 5 μm particle size; Macherey-Nagel) using a flow rate of 6 ml·min^-1^ (A: ACN with 1 % TFA, B: water with 1 % TFA). Obtained pure fractions were pooled and lyophilized. Peptide characterization was performed by analytical HPLC (1260 Infinity, Agilent Technology; flow rate of 1 ml·min^-1^, A: ACN with 1 % TFA, B: water with 1 % TFA) coupled with a mass spectrometer (6120 Quadrupole LC/MS, Agilent Technology) using electro spray ionization (Agilent Eclipse XDB-C18 culumn, 4.6×150 mm, 5 μm particle size). Analytical HPLC chromatograms were recorded at 210 nm **(Supplementary Figure 7)**. Quantification was performed by HPLC-based comparison (chromatogram at 210 nm) with a reference peptide **(Supplementary Table 2)**.

### Preparation of Recombinant Proteins

Arabidopsis PCO1 and PCO4 sequences in pDEST17 bacterial expression vectors (Invitrogen) were kindly provided by F. Licausi and J. van Dongen.^28^ Plasmids were transformed into BL21(DE3) *Escherichia coli* cells, and expression of recombinant protein carrying an N-terminal hexahistidine tag was induced with 0.5 mM IPTG and subsequent growth at 18°C for 18 hours. Harvested cells were lysed by sonication and proteins purified using Ni^++^ affinity chromatography, before buffer exchange into 250 mM NaCl/50 mM Tris-HCl (pH 7.5). Analysis by SDS-PAGE and denaturing liquid-chromatography mass spectrometry (LC-MS) showed proteins with more than 90% purity and with the predicted molecular weights.

The coding sequence of Arabidopsis ATE1 was cloned according to gene annotations at TAIR (www.arabidopsis.org) from cDNA. The sequence was flanked by an N-terminal tobacco etch virus (TEV) recognition sequence for facilitated downstream purification (“tev”:ENLYFQ-X) using the primers ate1_tev_ss (5’-GCTTAGAGAATCTTTATTTTCAGGGGATGTCTTTGAAAAACGATGCGAGT-3’) and ate1_as (5’-GGGGACCACTTTGTACAAGAAAGCTGGGTATCAGTTGATTTCATACACCATTCTCTC-3’). A second PCR using the primers adapter (5’-GGGGACAAGTTTGTACAAAAAAGCAGGCTTAGAGAATCTTTATTTTCAGGGG-3’) and ate1_as was performed to amplify the construct to use it in a BP reaction for cloning into pDONR201 (Invitrogen) followed by an LR reaction into the vector pDEST17 (Invitrogen). The N-terminal hexahistidine fusion was expressed in BL21-CodonPlus (DE3)-RIL *Escherichia coli (E. coli)* cells. The expression culture was induced with 1 mM IPTG at OD=0.6 and grown for 16 hours at 18°C. After resuspension in LEW buffer (50 mM NaH2PO4, pH 8; 300 mM NaCl; 1 mM DTT), the cells were lysed by incubation with 1.2 mg/ml lysozyme for 30 min and underwent subsequent sonification in the presence of 1 mM PMSF. Recombinant protein was purified by Ni++ affinity chromatography and subjected to Amicon Ultra-15 (30K) (Merck Millipore) filtration for buffer exchange to imidazole-free LEW containing 20% glycerol.

### PCO Activity Assays and MALDI Analysis

PCO activity assays were conducted under the following conditions unless otherwise stated: 1 μM PCO1 or 4 was mixed with 100 or 200 μM RAP2_2-11_ peptide in 250 mM NaCl, 1 mM dithiothreitol (DTT), 50 mM Tris-HCl pH 7.5 and incubated at 30°C for 30-60 minutes. Addition of exogenous Fe(II) and/or ascorbate were not required for activity. Assays were stopped by quenching 1 μL sample with 1 μL alpha-cyano-4-hydroxycinnamic acid (CHCA) matrix on a MALDI plate prior to product mass analysis using a Sciex 4800 TOF/TOF mass spectrometer (Applied Biosystems) operated in negative ion reflectron mode. The instrument parameters and data acquisition were controlled by 4000 Series Explorer software and data processing was completed using Data Explorer (Applied Biosystems).

To test the activity of PCO4 in the presence of ^18^O_2_, 100 μL of an anaerobic solution of 100 μM RAP2_2-11_ in 250 mM NaCl/50 mM Tris-HCl pH 7.5 was prepared in a septum-sealed glass vial by purging with 100% N_2_ for 10 minutes at 100 mL/min using a mass flow controller (Brooks Instruments), as used for previous preparation of anaerobic samples to determine enzyme dependence on O_2_.^24^ PCO4 was then added using a gas-tight Hamilton syringe, followed by purging with a balloon (approx. 0.7 L) of ^16^O_2_ or ^18^O_2_ over the course of 10 minutes at room temperature. Reaction vials were then transferred to 30°C for a further 20 minutes before products were analysed by MALDI-MS as described above.

PCO4 activity was additionally tested in the presence of H_2_^18^O by conducting an assay in 75% H_2_^18^O, 25% H_2_O (with all enzyme/substrate/buffer components comprising a portion of the H2O fraction). Assays were conducted for 10 minutes at room temperature followed by 20 minutes at 30°C for comparison with assays conducted with ^18^O_2_. Products were analysed by MALDI-MS, as described above.

### UPLC-MS and MS/MS Analysis of PCO Assay Products

Ultra-high performance chromatography (UPLC) mass spectrometry (MS) measurements were obtained using an Acquity UPLC system coupled to a Xevo G2-S Q-ToF mass spectrometer (Waters) operated in positive electrospray mode. Instrument parameters, data acquisition and data processing were controlled by Masslynx 4.1. Source conditions were adjusted to maximize sensitivity and minimize fragmentation while Lockspray was employed during analysis to maintain mass accuracy. 2 μL of each sample was injected on to a Chromolith Performance RP-18e 100-2 mm column (Merck) heated to 40°C and eluted using a gradient of 95 % deionized water supplemented with 0.1 % (v/v) formic acid (analytical grade) to 95 % acetonitrile (HPLC grade) and a flow rate of 0.3 mL/min. Fragmentation spectra of substrate and product peptide ions (MS/MS) were obtained using a targeted approach with a typical collision-induced dissociation (CID) energy ramp of 30 to 40 eV. Analysis was carried out with the same source settings, flow rate and column elution conditions as above.

### ^1^H NMR Assay

Reaction components (5 μM PCO1 or PCO4 and 500 μM RAP2_2-11_) were prepared to 75 μL in 156 mM NaCl, 31 mM Tris-HCl (pH 7.5) and 10% D_2_O (enzyme added last), in a 1.5 mL microcentrifuge tube before being transferred to a 2 mm diameter NMR tube. ^1^H NMR spectra at 310 K were recorded using a Bruker AVIII 600 (with inverse cryoprobe optimized for ^1^H observation and running topspin 2 software; Bruker) and reported in p.p.m. relative to D_2_O (δ_H_ 4.72). The deuterium signal was also used as internal lock signal and the solvent signal was suppressed by presaturating its resonance.

### Arginylation Assay

The conditions for arginylation of the 12-mer peptide substrates were modified from ^43^. In detail, ATE1 was incubated at 10 μM in the reaction mixture containing 50 mM HEPES, pH 7.5; 25 mM KCl; 15 mM MgCh; 1 mM DTT; 2.5 mM ATP; 0.6 mg/ml *E. coli* tRNA (R1753, Sigma); 0.04 mg/ml *E. coli* aminoacyl-tRNA synthetase (A3646, Sigma); 80 μM (4 nCi/μ1) ^14^C-arginine (MC1243, Hartmann Analytic); 50 μM C-terminally biotinylated 12-mer peptide substrate and, where indicated, 1 μM purified recombinant PCO1 or PCO4 in a total reaction volume of 50 μL. The reaction was conducted at 30°C for 16 to 40 hours. After incubation, each 50 μL of avidin agarose bead slurry (20219, Pierce) equilibrated in PBSN (PBS-Nonidet; 100 mM NaH_2_PO_4_; 150 mM NaCl; 0.1% Nonidet-P40) was added to the samples and mixed with an additional 350 μL of PBSN. After 2 hours of rotation at room temperature, the beads were washed 4 times in PBSN, resuspended in 4 mL of FilterSafe scintillation solution (Zinsser Analytic) and scintillation counting was performed using a Beckmann Coulter LS 6500 Multi-Purpose scintillation counter.

## Acknowledgements

Petra Majovsky, Domenika Thieme and Wolfgang Hoehenwarter from the Proteomics Unit of the Leibniz Institute of Plant Biochemistry (IPB), Halle, are acknowledged for mass spectrometry of recombinant ATE1. David Staunton from the Biophysical Facility, Department of Biochemistry, University of Oxford is acknowledged for multi-angle light scatter analysis of PCO1/4. Geoff Grime from the University of Surrey Ion Beam Centre is acknowledged for assistance with the microPIXE data collection. This work was supported by a Biotechnology and Biological Sciences Research Council (U.K.) New Investigator grant (BB/M024458/1) to E.F., a grant for setting up the junior research group of the *ScienceCampus Halle-Plant-based Bioeconomy* to N.D., by a Ph.D. fellowship of the Landesgraduiertenförderung Sachsen-Anhalt awarded to C.N., by an Engineering and Physical Sciences Research Council (U.K.) studentship (EP/G03706X/1) to J.B-B., a Royal Society Dorothy Hodgkin Fellowship to E.F., a William R. Miller Junior Research Fellowship (St. Edmund Hall, Oxford) to R.H. and grant DI 1794/3-1 by the German Research Foundation (Deutsche Forschungsgemeinschaft, DFG) to N.D. Financial support came from the Leibniz Association, the state of Saxony Anhalt, the Deutsche Forschungsgemeinschaft (DFG) Graduate Training Center GRK1026 “*Conformational Transitions in Macromolecular Interactions*” at Halle, and the Leibniz Institute of Plant Biochemistry (IPB) at Halle, Germany. We thank Prof. J. van Dongen (RWTH Aachen University, Germany) and Prof. F. Licausi (Scuolo Superiore Sant'Anna, Pisa, Italy) for sharing pDEST-PCO plasmids and helpful discussions.

## Author Contributions

M.W. performed the PCO1/4 activity assays and MALDI/LC-/MS/MS analyses. M.K. performed and established arginylation reactions on peptides coupled to biotin pulldown and scintillation measurements and purified ATE1 protein. R.H. performed the NMR assays with E.F. D.W. prepared the pDEST17-PCO1 and 4 plasmids. C.M. synthesized the peptides, T.N.G. supervised and designed the synthesis, C.N. cloned and established purification and activity assays for ATE1. R.O. conducted LC-MS to analyse +12 Da mass shifts. J.W. performed LC-MS analysis. J.Y. and J.C.B-B. prepared samples for micro-PIXE analysis and J.C.B-B. and E.F.G. collected and analysed micro-PIXE data. This work was supported by the network of the European Cooperation in Science and Technology (COST) Action BM1307—” European network to integrate research on intracellular proteolysis pathways in health and disease (PROTEOSTASIS). E.F. performed the PCO1 and PCO4 protein purification and selected activity assays. E.F., M.W., M.K. and N.D. designed the study, E.F. and N.D. wrote the manuscript. M.W., M.K., N.D. and E.F. designed the figures. All authors read and approved the final version of this manuscript.

